# Persistent DNA methylation and downregulation of homeostatic genes in astrocytes after pilocarpine-induced status epilepticus: Implications for epileptogenesis

**DOI:** 10.1101/2024.12.31.630767

**Authors:** Dante Gómez Cuautle, Alicia Rossi, Alejandro Villarreal, Luciana D’Alessio, Alberto Javier Ramos

## Abstract

Epilepsy is a debilitating neurological disorder characterized by recurrent seizures, affecting millions of patients worldwide. Retrospective studies in Temporal lobe epilepsy (TLE) patients have shown a high incidence of an initial precipitating event (IPE) in early childhood followed by a silent period where epileptogenesis occurs to end up in chronic epilepsy. Epileptogenesis, the process through which a normal brain undergoes structural and functional changes leading to epilepsy, remains an enigmatic phenomenon. We hypothesized that epigenetics may be involved in epileptogenesis and specifically astrocytes could be affected by pathological remodeling. To study this process, we used three approaches: The lithium-pilocarpine model of TLE in rats, primary astroglial cultures exposed to epileptogenic DAMP named HMGB1, and brain tissue samples resected from TLE patients with drug-resistant epilepsy. We found that the IPE achieved by lithium-pilocarpine treatment (127/30 mg/kg IP) induced the hypermethylation of astrocytes at 7-, 21-, and 35 days post-IPE, indicating persistent epigenetic alterations in astrocytes during the epileptogenic period. In addition, we observed the downregulation of homeostatic astroglial genes AQP4; glutamine synthase (GS), and Kir4.1 with increased proinflammatory genes (C3, MAFG) and DNA methyl transferases (DNMT) expression. These alterations were mimicked in primary astrocyte cultures exposed to the epileptogenic HMGB1 (500 ng/ml; 18 hours) that also induced the hypermethylation of homeostatic astroglial genes. Astrocytes from TLE patients brains showed reactive astrogliosis, increased DNA methylation, and downregulation of homeostatic genes Kir4.1 and GS. These findings show that astrocytes are pathologically altered during the epileptogenic period, combining the proinflammatory gain of function with the loss of homeostatic profile. This may sustain the long-term alterations underlining epileptogenesis.

## Introduction

Epilepsy is a chronic neurological disorder that affects almost 60 million people worldwide (McNamara et al., 2006) and is characterized by a propensity for spontaneous, recurrent seizures, arising from aberrant neuronal activity that produces an excitatory/inhibitory disbalance (Fisher et al., 2014). While anti-seizure medications provide symptomatic relief to many patients, they fail to address the fundamental processes driving epileptogenesis, leading to drug resistance in a significant proportion of cases (Clossen and Reddy, 2017; Lazarowski & Czornyj, 2013; Łukasiuk and Lason, 2023). Understanding the pathophysiology of epileptogenesis is crucial for developing targeted interventions to prevent or halt the development of epilepsy and going beyond the mere treatment of the symptoms.

Epileptogenesis involves complex cellular and molecular changes in the brain that include neuronal alterations, spurious synaptogenesis, reactive gliosis, neuroinflammation, and other alterations leading to establishing a hyperexcitable network. Among these changes, reactive gliosis and neuroinflammation have emerged as prominent features (Binder et al., 2004; Rossi et al., 2013; 2017; Vezzani et al., 2022). Astrocytes, the most abundant glial cells in the brain, respond to insults such as seizures, traumatic brain injury, or inflammation by undergoing reactive gliosis. In this state, astrocytes undergo morphological and functional changes, releasing proinflammatory cytokines, chemokines, and growth factors that can modulate synaptic activity and neuronal excitability (Verkhratsky et al., 2017; Vargas-Sánchez et al., 2019). Recent evidence from experimental models and transcriptomic studies has shown that reactive gliosis may be detrimental to neuronal survival. In this situation, the astrocytes engage in pathological remodeling with overexpression of proinflammatory factors and downregulation of homeostatic genes required for nervous system homeostasis, neuronal survival, and synaptic maintenance (Verkhratsky et al 2017, 2021; Escartin et al., 2021; Cieri et al., 2023; Gomez Cuautle et al., 2024).

Epigenetic modifications, including DNA methylation, histone modifications, and non-coding RNA regulation, have garnered increasing attention in the context of epilepsy (Miller-Delaney et al., 2015; Williams-Karnesky et al., 2013; Zhang et al., 2021). DNA methylation, the addition of a methyl group to cytosine residues within CpG dinucleotides, is one of the most extensively studied epigenetic marks. DNA methylation patterns are dynamically regulated during brain development and adult neuroplasticity, affecting gene expression and influencing neuronal function (Hsieh and Eisch, 2010). Dysregulation of DNA methylation has been observed in various neurological disorders, and accumulating evidence points to its potential role in epileptogenesis. Particularly in reactive astrogliosis, we have shown that lipopolysaccharide (LPS) exposure, and traumatic or ischemic brain injury activate epigenetic remodeling mechanisms in astrocytes (Villarreal et al., 2021; Gomez Cuautle et al., 2024). While histone acetylation seems to be involved in the proinflammatory gene expression, DNA methylation seems to be affecting the expression of astroglial genes involved in homeostasis (Gomez Cuautle et al., 2024; Wheeler et al., 2023).

The role of DNA methylation in epilepsy has been proposed, although initially focusing on neurons (Williams-Karnesky et al., 2013; Miller-Delaney et al., 2015; Tao et al., 2021; Martins-Ferreira et al., 2022). However, it has been suggested that there is a link between reactive gliosis and epigenetic mechanisms, particularly DNA methylation, in epilepsy (Robel et al., 2015; Berger et al., 2019; Çarçak et al., 2023). Boni and colleagues (2020) demonstrated that Kir4.1 K^+^ rectifying channel expression in astrocytes is controlled by DNA methylation. Berger and colleagues (2020) also proposed that Cox2 and Cxcl10 proinflammatory molecules are controlled by DNA methylation in astrocytes. Based on this evidence and epigenetic mechanisms’ proposed role in epileptogenesis, we hypothesized that epigenetic remodeling and astroglial pathological conversion occur during the epileptogenic process, leading to pathologically remodeled astrocytes that are not fully capable of sustaining neuronal homeostasis. The previous evidence is probably the *tip of the iceberg* of a more profound pathological gene expression program occurring in astrocytes during the epileptogenic process. To test this, we here studied the astroglial alterations during the epileptogenic latency period that follows a pilocarpine-induced status epilepticus and analyzed the cellular pathways using primary hippocampal astroglial cultures. The long-term status of these modifications in astrocytes was confirmed on human brain samples resected from drug-resistant TLE patients.

## Materials and methods

### Experimental animals

All procedures involving animals and their care adhered to our institutional guidelines, which align with the NIH guidelines for the Care and Use of Laboratory Animals and the principles outlined in the Guidelines for the Use of Animals in Neuroscience Research by the Society for Neuroscience. These procedures were approved by the CICUAL committee of the School of Medicine, University of Buenos Aires (Res. Nr. 2367/2018 and 74747/2019). Every effort was made to minimize animal suffering and reduce the number of animals used. The animal treatment, surgeries, and euthanasia were performed at the Animal Facility of the IBCN, School of Medicine, University of Buenos Aires.

### Human samples

Hippocampal and temporal-pole cortex tissue sections obtained from surgical pieces collected from patients who underwent epilepsy surgery (anterior temporal lobectomy) for drug resistant TLE within the period from 2006 to 2016 year, were selected. Also, postmortem brain sections from controls matched by age and sex were included. Clinical, neurological, and pathological data from these cohorts was previously reported (D’Alessio et al., 2015, 2020). All patients were enrolled in a clinical protocol for epilepsy surgery approved by the Ethics Committee of Ramos Mejía Hospital and all of them signed the informed consent for participation. All patients were receiving their habitual medication at the moment of epilepsy surgery and had hippocampal sclerosis with long-term seizure events before surgery (D’Alessio et al., 2015; 2020).

### Data availability

All primary data are available from authors upon reasonable request. Further experimental details are also available from authors upon reasonable request.

### Materials

Primary antibodies from different brands were used as follows: rabbit polyclonal anti-glutamine synthetase (ThermoFisher, RRID:AB_2546416; 1/500), chicken polyclonal anti-Glial Fibrillary Acidic Protein (GFAP) (ThermoFisher, RRID:AB_1074620; 1/500), mouse monoclonal anti-GFAP (Sigma, RRID:AB_477010; 1/1000), rabbit polyclonal anti-GFAP (Dako, RRID:AB_10013382; 1:3000), rabbit polyclonal anti-AQP4 (Santa Cruz, RRID:AB_2274338; 1/800); mouse monoclonal anti-5-methylcytosine (5mC) (Zymo Research cat# A-3001, 1/1000). Secondary fluorescent antibodies were obtained from Jackson Immunoresearch and Sigma. Cell culture reagents were obtained from ThermoFisher (Carlsbad, USA), and fetal calf serum (FCS) was purchased from Natocor (Córdoba, Argentina). Poly-L-lysine (Sigma, cat# P4707), DAPI (4’,6-diamidino-2- phenylindole; Sigma, cat# D9542), HMGB1 (human recombinant Sigma, cat# H4652), and other chemicals were obtained from Sigma (USA).

### Animal Procedures

Adult male Wistar rats (200–250 g) from the animal breeding facility of the IBCN, Facultad de Medicina (University of Buenos Aires) were used in this study. Animals were housed under controlled conditions of temperature, humidity, and lighting (18–25 °C, 60% humidity, 12 h/12 h light/dark cycle) with standard laboratory rat food and water ad libitum, under the continuous supervision of professional technicians. Rats were subjected to the lithium-pilocarpine model of Temporal Lobe Epilepsy (TLE) following the protocols outlined by Rossi et al. (2013) and Sarchi et al. (2023). Briefly, animals received an intraperitoneal injection of 3 mEq/kg lithium chloride and were randomly assigned to one of two experimental groups. After 20 hours, one group was administered 30 mg/kg pilocarpine, while the other received a saline injection. Seizure behavior was monitored using the Racine scale, with Status Epilepticus (SE) defined as sustained seizures scoring between 3 and 5 on the Racine scale for a minimum of 5 minutes without reverting to lower stages. Approximately 70% of the pilocarpine-treated rats developed SE within an hour (SE group), while the remaining 30% displayed mild seizure activity corresponding to Racine stages 1–2 (No SE group). To minimize mortality, seizures were terminated after 20 minutes using an intraperitoneal injection of 20 mg/kg diazepam. Animals were euthanized at 7-, 21-, or 35-days post-SE (DPSE) under deep anesthesia with ketamine/xylazine (90/10 mg/kg, i.p.).

### Primary astroglial cultures

Neonatal Wistar rats (3 days old; 6–8 pups per experiment) were used, following the protocols of Rosciszewski et al. (2019) and Villarreal et al. (2021). Briefly, pups were rapidly decapitated, and their brains were immediately placed in cold Dulbecco’s Modified Eagle Medium (DMEM; Invitrogen, cat# 119965092). Under a microscope, the meninges were carefully removed, and the hippocampi were dissected and mechanically dissociated using tweezers. A second round of dissociation was performed by pipetting with progressively smaller micropipette tips until no visible cell clumps remained. The resulting cell suspension was transferred to 15 ml conical tubes and centrifuged at 1000 RPM for 5 minutes. The supernatant was discarded, and the cell pellet was resuspended in DMEM supplemented with 10% fetal calf serum, 2 mM L-glutamine, and 100 μg/ml penicillin/streptomycin (Sigma, cat# P4333). Cells were plated in poly-L-coated flasks and incubated at 37°C in a 5% CO_2_ atmosphere until reaching confluence, typically within 7–10 days. The medium was changed one day after plating and again on day three. Confluent cultures were prepared either as mixed glial cultures or microglia-depleted astrocyte-enriched cultures. Untreated flasks yielded mixed glial cultures containing approximately 20% microglial cells, as previously described (Rosciszewski et al., 2018, 2019; Villarreal et al., 2021). To generate astrocyte-enriched cultures, confluent mixed cultures were subjected to orbital shaking at 140 RPM for 24 hours, followed by washing with pre-warmed supplemented DMEM and treatment with 50 μg/ml 5-fluorouracil for 24 hours to inhibit mitotic microglia (Rosciszewski et al., 2018). Cultures were then maintained in supplemented DMEM until re-plating for experiments. For experimental procedures, cells from either mixed glial or enriched astrocyte cultures were detached using 0.05% trypsin (Sigma, cat# T6567) and re-plated onto poly-L-coated multiwell plates. Cultures were treated with HMGB-1 (500 ng/ml) or vehicle, as described by Rosciszewski et al. (2019). After experimental treatments, cells were washed with PBS (pH 7.2) and fixed with 4% paraformaldehyde and 4% sucrose in PBS for 15 minutes at room temperature. Fixed cells were then washed three times with PBS and stored at 4°C until immunofluorescence staining.

### Immunocytochemistry

For immunocytochemistry, fixed cells were permeabilized by treatment with 0.1% Triton-X in PBS for 15 minutes. For 5mC immunostaining, additional acidic treatment was applied: fixed and permeabilized cells were exposed to 2N HCl for 1 hour at room temperature, followed by neutralization with 0.1 M Tris-HCl (pH 8.4). Cells were then incubated with a blocking solution (5% normal horse serum in PBS) for 30 minutes at room temperature to prevent nonspecific binding. Subsequently, cells were incubated overnight at 4°C with primary antibodies diluted in the blocking solution. After this period, unbound primary antibodies were removed by washing the cells three times with PBS. Visualization of target antigens was achieved by incubating the cells with fluorescently labeled secondary antibodies for 4 hours at room temperature. Finally, a nuclear counterstain was performed using 0.1 μg/ml DAPI. Similarly, brain sections were processed in a free-floating state as described by Aviles Reyes et al. (2009) and Sarchi et al. (2023). Epifluorescence images were captured using an Olympus IX-81 microscope with a DP71 camera (Olympus, Japan) or a Zeiss LSM-880 confocal microscope equipped with an Airyscan module (Carl Zeiss, Germany; Hoff, 2015). Details of the secondary antibodies used are provided in Table 3.

### Real-time RT-PCR

For real-time reverse transcription polymerase chain reaction (RT-PCR) analysis, astroglial-enriched cultures were rapidly lysed directly in the culture plates, and RNA was extracted using the Zymo Quick-RNA Miniprep Kit x 50 (Zymo, USA; cat# ZRR1054), following the manufacturer protocol. Reverse transcription was performed using 200 ng of total RNA and the MMLV RT kit (Genbiotech) to synthesize cDNA. Quantitative PCR (qPCR) was conducted on an AriaMx Real-Time PCR System (Agilent, USA) using 5x HOT FIREPol EvaGreen qPCR Mix Plus Rox (Solis Biodyne, cat# 08-24-00001) and 5 μM of each primer. The qPCR thermal profile consisted of an initial denaturation at 95°C for 2 minutes, followed by 40 cycles of 15 seconds at 95°C, 15 seconds at 60°C, and 60 seconds at 72°C, with a final melting curve analysis. Relative RNA expression was determined using the change-in-threshold (2−ΔΔCT) method, normalizing the data to the housekeeping gene TATA box binding protein (TBP). For the heatmap data, the ΔΔCt results were transformed using a base-2 logarithm, centering the values around the control to facilitate comparison between experimental groups. The PCR products were also analyzed on a 2% agarose gel, and images were captured using a gel documentation system (Gel Doc™ EZ System, Bio-Rad). Specific primer sequences are detailed in Table 1. All qPCR reactions were performed in triplicate. Data analysis was carried out using GraphPad Prism version 9 for Windows (GraphPad Software, La Jolla, California, USA).

**TABLE 1:**
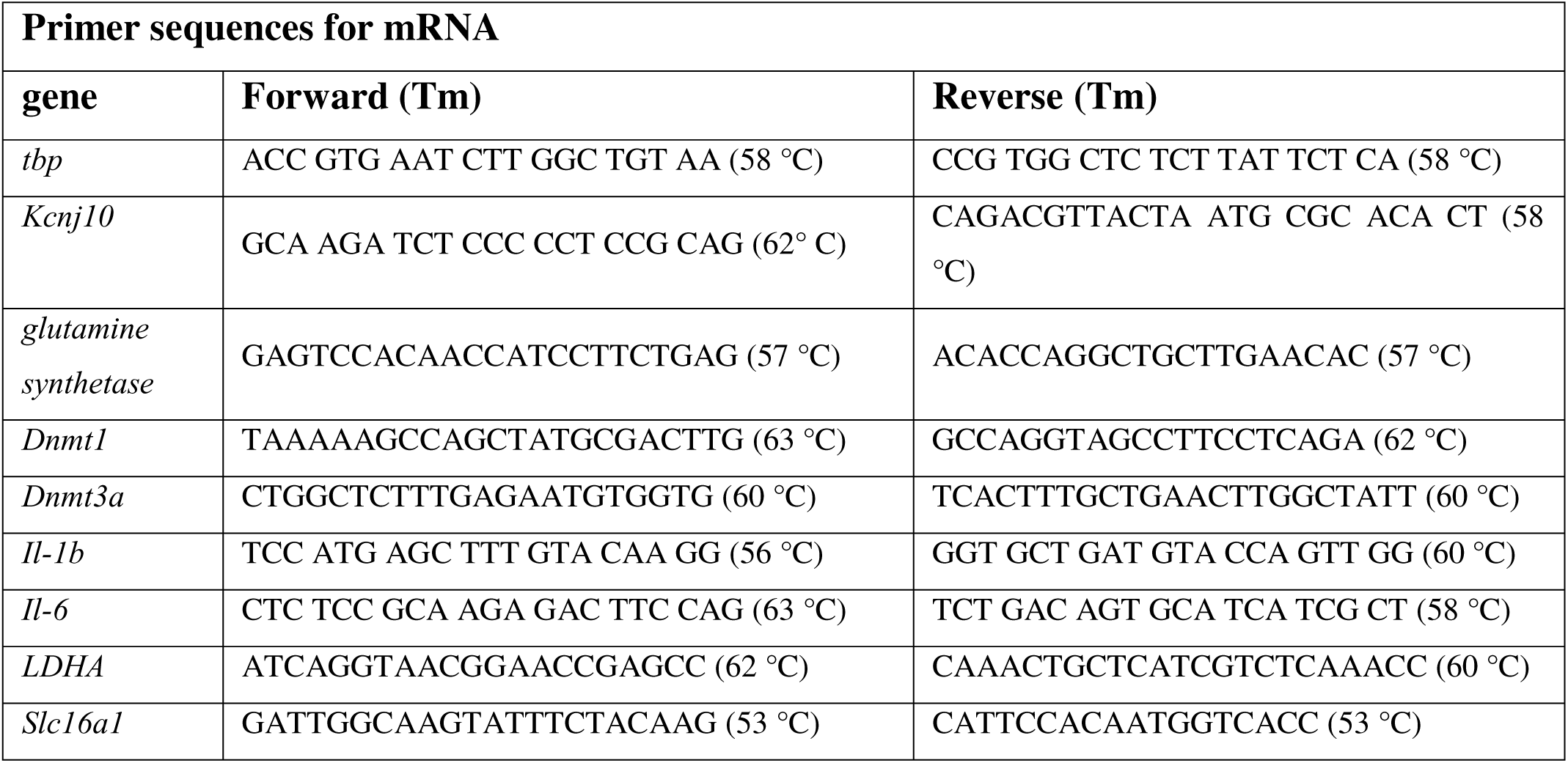
Sequences of primers for mRNA.

### Methylation-sensitive PCR

Genomic DNA was extracted from primary astroglial cultures using the PuriPrep-T Kit (Inbio Highway, Argentina) according to the manufacturer’s instructions. DNA concentration and purity were assessed with a Nanodrop® spectrophotometer (Thermo Fisher Scientific, Wilmington, DE, USA). Subsequently, 500 ng of the extracted genomic DNA underwent bisulfite conversion using the EZ DNA Methylation™ Kit (Zymo Research, Irvine, CA, USA), following the manufacturer’s protocol. Primers for methylation-specific PCR (Table 2) were designed to flank CG-rich regions identified through the UCSC Genome Browser using the Bisulfite Primer Seeker tool (www.zymoresearch.com/tools/bisulfite-primer-seeker). The promoter region sequences were: **Slc16a1** (RGSC 6.0/rn6 chr2:207108143-207109238, 1096 bp), **LDHA** (RGSC 6.0/rn6 chr6:33407166-33407450, 285 bp), and **GS** (RGSC 6.0/rn6 chr13:71330754-71331828, 1075 bp). Methylation-specific PCR was performed using the Zymo EZ DNA Methylation-Direct Kit (cat# D5020) per the manufacturer instructions. In brief, astrocyte-enriched cultures were lysed directly in culture plates and treated with proteinase K at 50°C for 20 minutes. The supernatant was transferred to PCR tubes, denatured at 98°C for 8 minutes, and incubated at 64°C for 3.5 hours in a thermal cycler. The samples were then transferred to columns, treated with a desulphonation reagent, washed and eluted for downstream PCR assays. The PCR reaction mixture (20 μl total volume) included 4 μl of 5x HOT FIREPol EvaGreen qPCR Mix Plus Rox™ polymerase, 0.5 μl each of forward and reverse primers, 3 μl (100 ng) of bisulfite-converted DNA, and PCR-grade H O to complete the volume. Reactions were run on an AriaMx Real-Time PCR System (Agilent) with the following thermal profile: initial denaturation at 95°C for 10 minutes; 40 cycles of 95°C for 50 seconds, annealing at a specific temperature for 45 seconds, and extension at 72°C for 60 seconds; followed by a final extension at 72°C for 5 minutes. PCR products were analyzed on 2% agarose gels, and images were captured using a Gel Doc™ EZ System (Bio-Rad). Quantitative analysis was performed with ImageJ-Fiji. To optimize resource use and minimize toxic waste (e.g., ethidium bromide), different lanes were used for separate samples. For clarity in figures, images were obtained from the same gel, cropped, and assembled. The ratio of methylated/unmethylated integrated density was calculated.

**TABLE 2:**
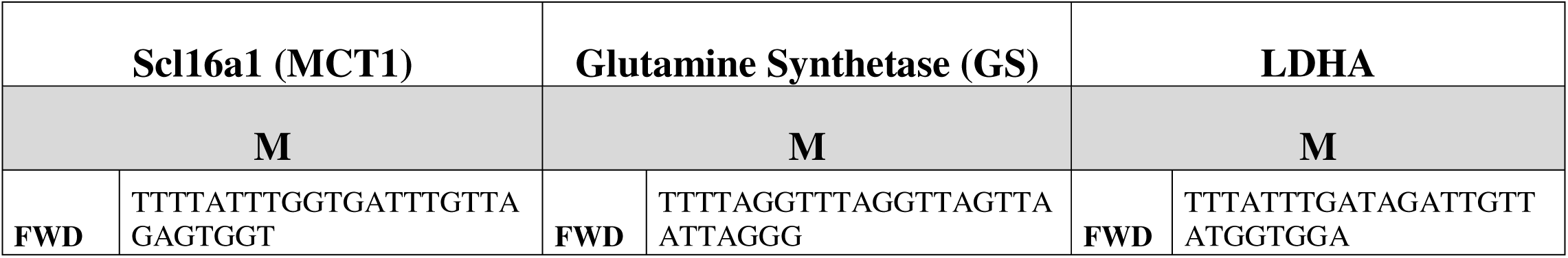
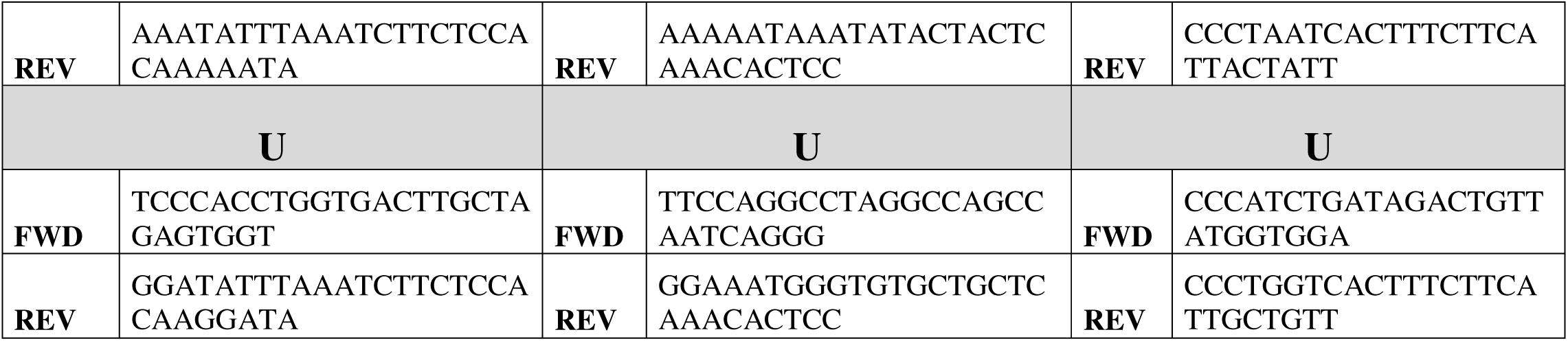
Sequences for methylation-sensitive PCR. Primers for methylated DNA (M primer set) and unmethylated DNA (U primer set) of the promoter regions were designed using the Bisulfite Primer Seeker software.

**TABLE 3:**
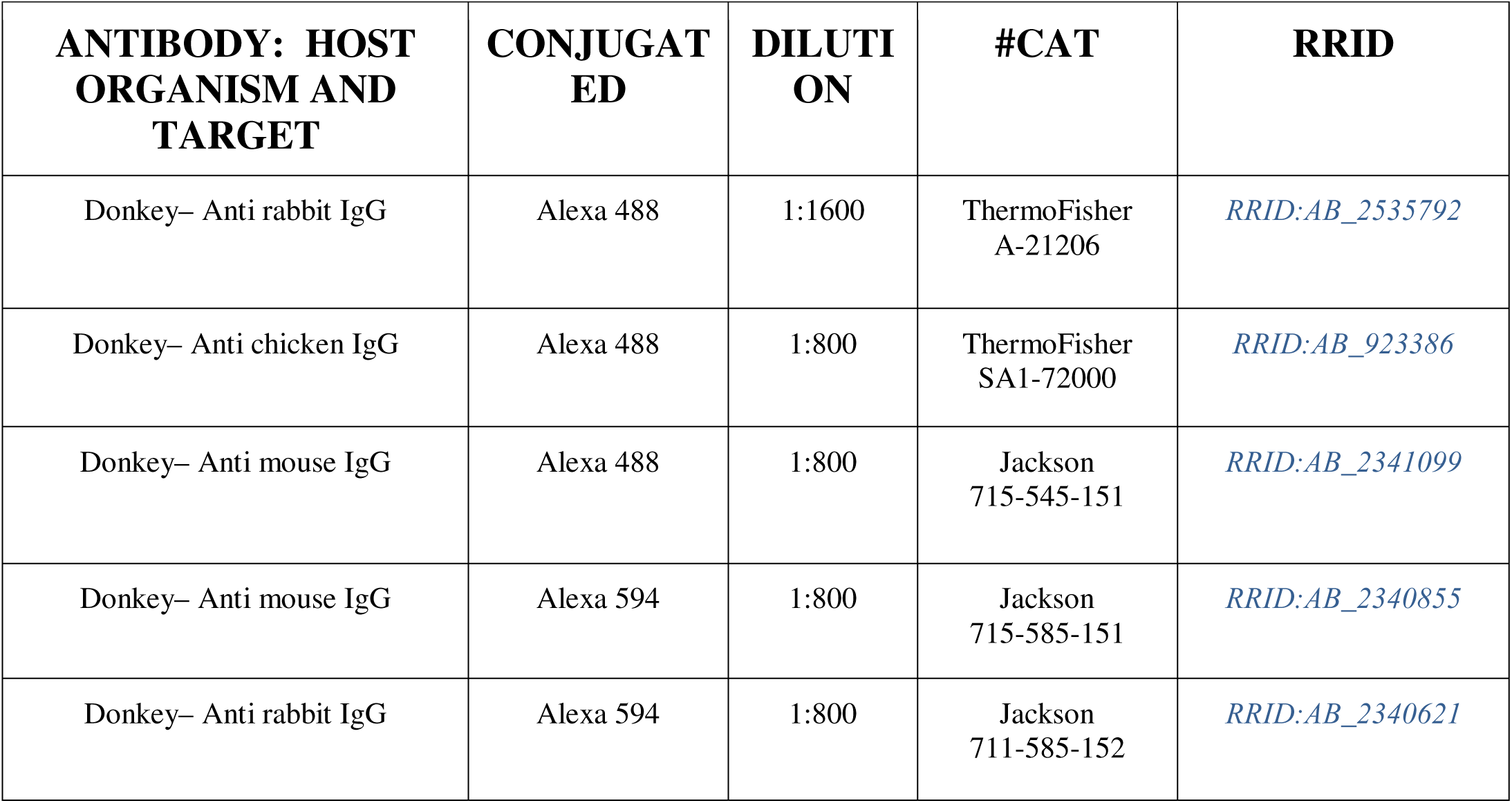
Secondary antibodies details.

| **ANTIBODY: HOST ORGANISM AND TARGET** | **CONJUGATED** | **DILUTION** | **#CAT** | **RRID** |
| --- | --- | --- | --- | --- |
| Donkey– Anti rabbit IgG | Alexa 488 | 1:1600 | ThermoFisher A-21206 | *RRID:AB_2535792* |
| Donkey– Anti chicken IgG | Alexa 488 | 1:800 | ThermoFisher SA1-72000 | *RRID:AB_923386* |
| Donkey– Anti mouse IgG | Alexa 488 | 1:800 | Jackson 715-545-151 | *RRID:AB_2341099* |
| Donkey– Anti mouse IgG | Alexa 594 | 1:800 | Jackson 715-585-151 | *RRID:AB_2340855* |
| Donkey– Anti rabbit IgG | Alexa 594 | 1:800 | Jackson 711-585-152 | *RRID:AB_2340621* |

## Results

### 1. Astroglial DNA methylation increases during the latency period that follows status epilepticus

Pilocarpine-induced status epilepticus (SE) is an animal model of IPE known to produce profuse reactive gliosis accompanied by significative neuronal degeneration as was previously reported (Rossi et al., 2013; 2017, Sarchi et al., 2023). These alterations persist during the latency period that follows pilocarpine-induced SE. Considering that 5mC abundance is altered in neurons after SE (Kobow and Blümcke, 2018; Tao et al., 2021), we investigated whether GFAP+ reactive astrocytes induced by SE also show alterations in their DNA methylation levels. As shown in Figure 1A-B-C, significative reactive astrogliosis was observed in the hippocampus starting at 7DPSE (DPSE, days post-status epilepticus) together with an increased abundance of 5mC immunostained glial cells that are observed in the stratum radiatum of CA1 and CA2-3 hippocampal regions. Specifically in hippocampal stratum radiatum, an area devoid of neuronal somas with mostly apical dendrites of pyramidal neurons, GFAP+ astrocytes showed an increase in the 5mC nuclei immunostaining in SE-exposed animals (Figure 1D). The hypermethylation is still significantly different from control animals at 21 and 35DPSE (Figure 1E). Detailed high-magnification images at 35DPSE showed that astroglial DNA is still hypermethylated at 35DPSE (Figure 1D) as the quantitative analysis showed (Figure 1E). These findings suggest that pilocarpine-induced SE promotes long-lasting DNA hypermethylation of reactive astrocytes.

**Figure 1:**
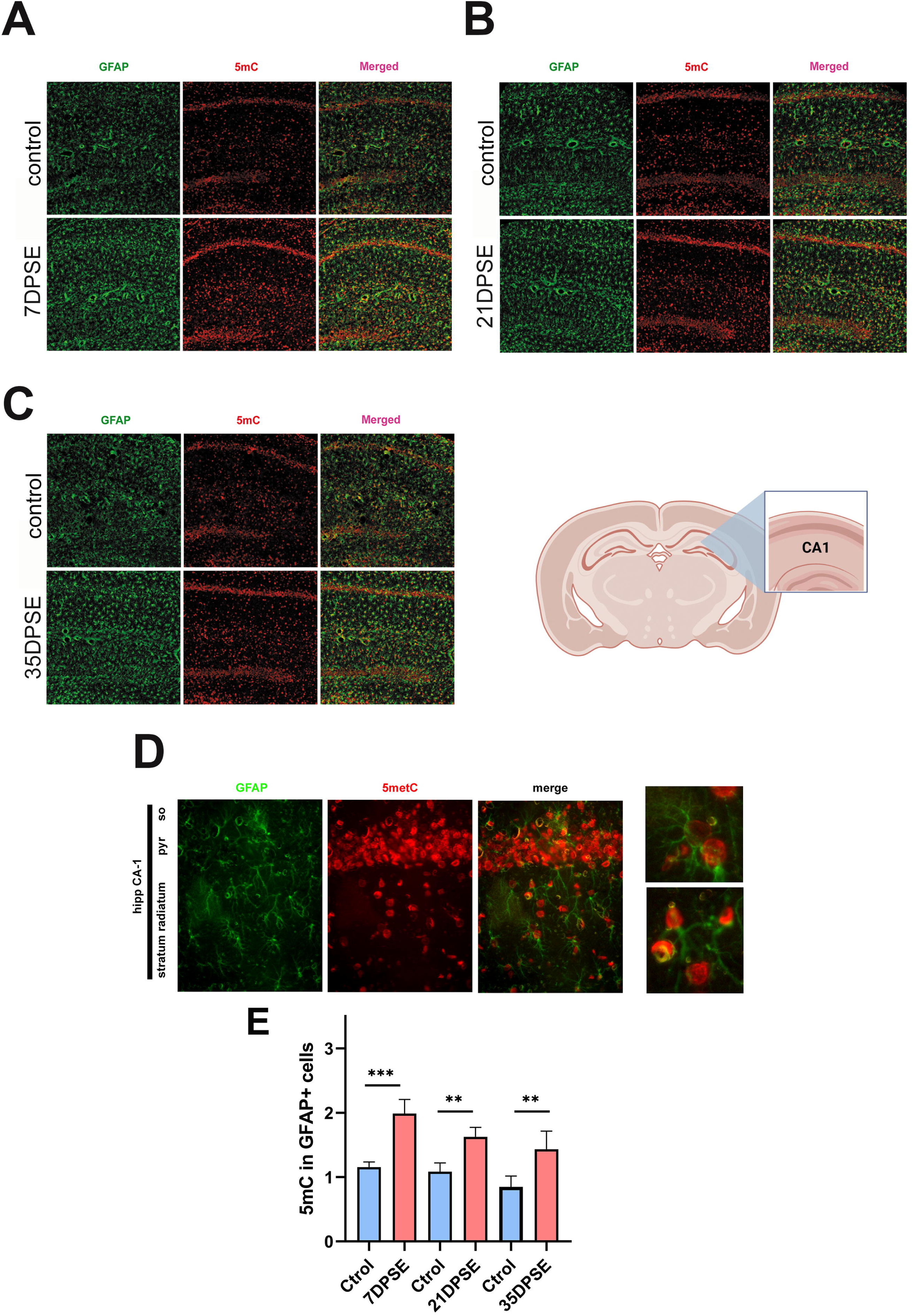
Increased DNA methylation in astrocytes after pilocarpine-induced SE. **A- B-C**: Representative images of 5mC/GFAP immunostaining in hippocampal brain sections of animals exposed to pilocarpine-induced SE at 7 days post-SE (DPSE) (A), 21DPSE (B) or 35DPSE (C). **D**: High magnification images showing details of the CA-1 area and astrocytes immunolabeled by 5mC at 35DPSE. **E**: Quantitative analysis of the 5mC abundance in GFAP+ cells as mean fluorescence intensity (MFI). Data are presented as mean ± SEM and significance was analyzed by two-way ANOVA and Tukey post-test. N= 5-7 animals per group.

### 2. Augmented expression of proinflammatory genes and reduced expression of astroglial homeostatic genes after status epilepticus

The pilocarpine-induced SE is known to produce neurodegenerative consequences during the latency period. Consistent with the hypothesis of the activation of a proinflammatory cascade in astrocytes, we observed that the proinflammatory-neurodegenerative astroglial marker C3 showed a significative increased expression after pilocarpine-induced SE at 7DPSE and 21DPSE (Figure 2A). Then, C3 expression showed a decrease to be normalized at 35DPSE (Figure 2A). On the other hand, the expression of the homeostatic K+ inward rectifying channel Kir4.1 (kcnj10 gene), a known gene target deeply affected by the DNA methylation in epilepsy (Boni et al., 2020) showed reduced expression in the latency period after SE (Figure 2B). The expression of GFAP and the astroglial neurotoxic factor and DNA methylation factor named MAFG were also increased after pilocarpine-induced SE (Figure 2C). Interestingly, and in agreement with the increased astroglial DNA methylation observed in Figure 1, the expression of both DNA methyl transferases DNMT1a and DNMT3a were also increased after pilocarpine-induced SE (Figure 2C). Having observed this proinflammatory and DNA methylation burst in astrocytes early after pilocarpine-induced SE, we analyzed whether the expression of classical astroglial homeostatic genes was altered by the pilocarpine-induced SE. As shown in Figure 2D and 2E immunohistochemistry showed that the homeostatic proteins AQP4 water channel (Figure 2D) and glutamine synthase (Figure 2E) were all decreased after pilocarpine-induced SE. We conclude that pilocarpine-induced SE produces a rapid activation of astroglial proinflammatory-neurodegenerative genes together with a long-lasting reduction in the expression of astroglial genes involved in CNS homeostasis like AQP4 and GS.

**Figure 2:**
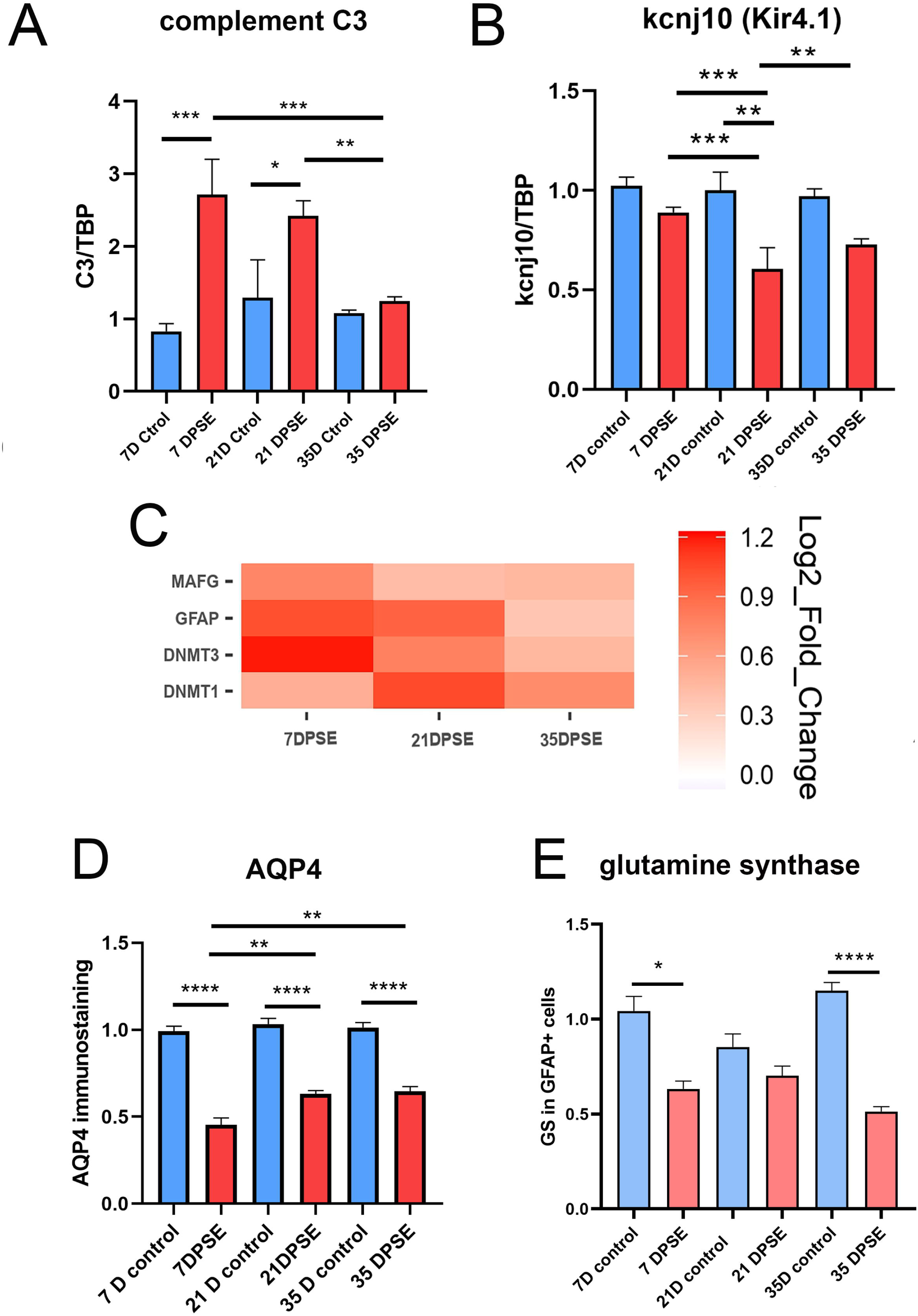
Increased proinflammatory gene expression together with reduced homeostatic genes in astrocytes after pilocarpine-induced SE. **A**: RT-PCR studies showing the expression of complement C3 mRNA relative to the housekeeping TBP gene in the hippocampus of animals exposed to pilocarpine-induced SE and control animals. Results are presented as mean ± SE and statistical analysis was performed with two-way ANOVA and Tukey post-test. N= 3-6 animals per group. **B:** RT-PCR studies showing the expression of kcnj10 (Kir 4.1 gene) mRNA relative to the housekeeping TBP gene in the hippocampus of animals exposed to pilocarpine-induced SE and control animals. Results are presented as mean ± SE and statistical analysis was performed with two-way ANOVA and Tukey post-test. N= 4-7 animals per group. **C**: The heat map illustrates the genes that were either downregulated or overexpressed following pilocarpine-induced SE at different time points after SE (DPSE) in the hippocampus. The values were log2-transformed after applying the ΔΔCt method to facilitate comparison between experimental and control groups. **D-E**: Quantitative analysis of AQP4 immunostaining (D) or glutamine synthase (GS) immunostaining in the hippocampus of animals exposed to pilocarpine-induced SE. Results are presented as mean ± SE and statistical analysis was performed with two-way ANOVA and Tukey post-test. N= 4-7.

### 3. DAMP HMGB1 exposure in primary astrocytes in culture induces hypermethylation, reduces expression of homeostatic genes, and increases proinflammatory gene expression

We next asked about the possible mechanistic explanation for the above-mentioned findings after pilocarpine-induced SE in rodents. Having in mind that HMGB1 is a DAMP released by degenerating neurons after SE (Ravizza et al., 2018; Vezzani et al., 2022) and seems to underlie the pro-epileptogenic and proinflammatory processes following SE (Maroso et al., 2010; Rosciszewski et al., 2019), we hypothesized that HMGB1 exposure may recapitulate in astrocytes the observations found in pilocarpine-exposed animals. To test this hypothesis, primary astroglial-enriched cultures were exposed to 500 ng/ml HMGB1 for 18 h, and cultures were followed for 24 h, 72 h, or 7 days after exposure. As shown in Figure 3A, astroglial DNA methylation assessed by the 5mC content was significantly increased after HMGB1 exposure. Interestingly, astroglial hypermethylation was steadily found to increase in the long term, persisting at 72 h and even after 7 days (Figure 3A). The observed long-lasting hypermethylation in astrocytes showed a correlation in the increased expression of DNA methyl transferases involved in DNA methylation named DNMT1 and DNM3a (Figure 3B-C). In addition, decreased expression of homeostatic genes kcnj10 (Kir 4.1), glutamine synthase (GS), Scl16a1 (MCT-1), and LDHA was also observed after the HMGB1 exposure (Figure 3B-C). Interestingly, the decrease in homeostatic gene expression in astrocytes showed an inverse correlation with proinflammatory genes IL-1B and IL-6 which were both augmented after HMGB1 and persisted increased for the 7 days of the observation period (Figure 3B-C). Specifically, IL-6 mRNA was persistently high, while IL-1B showed a trend to be reduced until 7 days after HMGB1 exposure. We conclude that 18h exposure to HMGB1 induced DNA hypermethylation in astrocytes together with increased proinflammatory gene expression and decreased levels of homeostatic gene expression.

**Figure 3:**
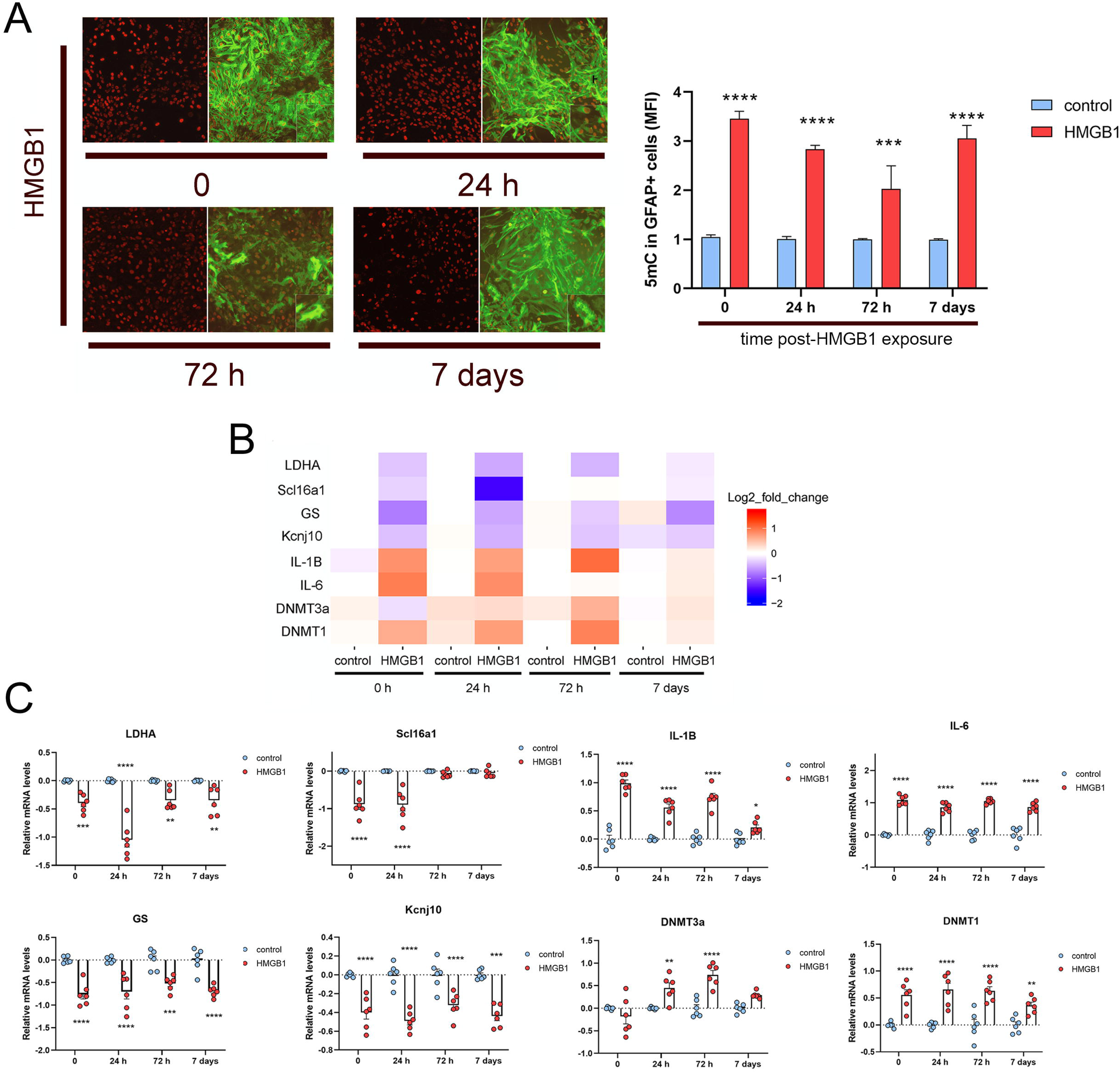
HMGB1 exposure induces astroglial hypermethylation, neuroinflammatory gene expression, and down-regulation of homeostatic genes. **A**: Astroglial cultures were exposed to HMGB1 500 ng/ml for 18 h and allowed to recover in the control tissue culture medium for 0, 24 h, 72 h, or 7 days. Representative images of the 5mC/GFAP immunostaining after the indicated times. The graph shows the quantitative analysis of the mean fluorescence intensity (MFI) in GFAP-immunoreactive cells in the culture relative to the control conditions. Results are presented as mean ± SE and statistical analysis was performed with two-way ANOVA and Tukey post-test. N= 9. **B**: The heat map illustrates the gene expression levels analyzed by real time-RT-qPCR at the different recovery time points after HMGB1 exposure. Data were log2-transformed after applying the ΔΔCt method to facilitate comparison. **C**: Detailed relative mRNA levels of the results presented in the heat map with the statistical analysis. Results are presented as mean ± SE and statistical analysis was performed with two-way ANOVA and Tukey post-test. N= 6.

### 4. Increased methylation of homeostatic gene promoters in astrocytes after HMGB1 exposure

The observed hypermethylation in astrocytes after pilocarpine-induced SE in rodents, added to the fact that HMGB1 induces astroglial hypermethylation in primary astrocytes in culture together with the repression of homeostatic genes and over-expression of proinflammatory genes; pointed us to analyze if specific gene promoters were hypermethylated after HMGB1 exposure. For this purpose, astrocytes were exposed to HMGB1 for 18 h, and methylation-sensitive PCR studies were performed on the previously described highly conserved sequences of these homeostatic gene promoters (Gomez Cuautle et al., 2024). As shown in Figure 4A, increased methylation of GS, Scl16a1 (MCT1), and LDH astroglial homeostatic genes were observed after HMGB1 treatment by the methylation-sensitive PCR assay. Quantitative studies showed significative differences as presented in Figure 4B. We conclude that these homeostatic genes show increased methylation after HMGB1 exposure and this may justify the reduced expression observed under these conditions.

**Figure 4:**
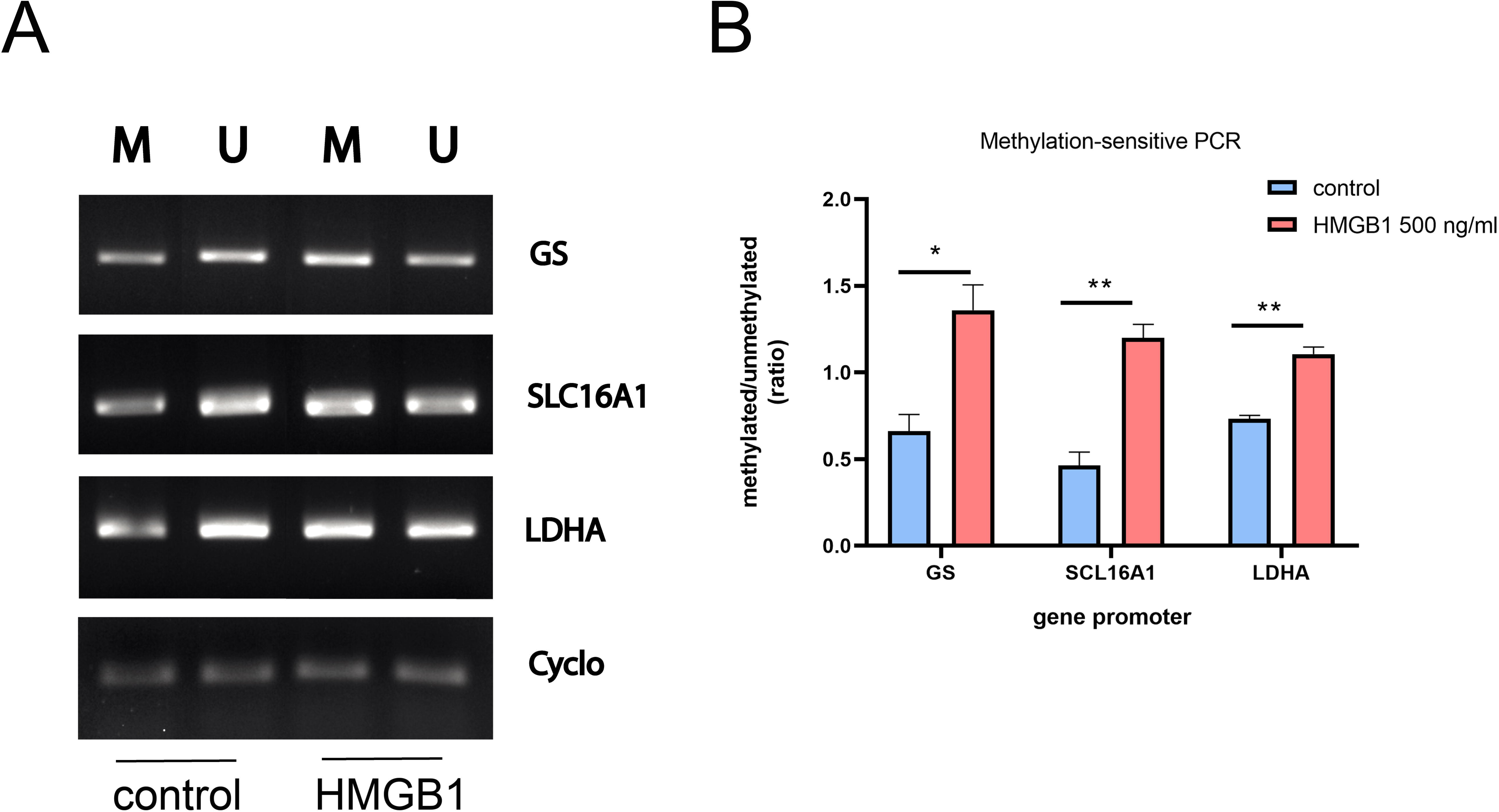
Increased methylation of homeostatic genes after HMGB1 exposure. **A**: Astroglial cultures were exposed to HMGB1 500 ng/ml for 18 h and recovered in the control tissue culture medium for 24 h. The highly conserved regions in the homeostatic gene promoters GS, scl16a1 (MCT-1), and LDHA were located using the UCSC genome browser based on rat DNA sequences as depicted in Gomez Cuautle et al., 2024. The CpG- rich islands were identified in silico using the Bisulfite Primer Seeker software (Zymo). After astroglial DNA purification and processing as explained in the methods section, PCR assays using primers for methylated and unmethylated sequences were run and products were analyzed and photographed in 2% agarose gels. A representative image of the 2% agarose gel is presented in this figure. **B**: Quantitative analysis was performed and the integrated density of the bands is shown in this figure as a ratio of the methylated/unmethylated integrated density. Results are presented as mean ± SEM and statistical analysis was performed with an unpaired t-test with Welch’s correction. N = 3.

### 5. Human TLE samples show reactive astrogliosis, increased DNA methylation, and reduced expression of homeostatic proteins Kir4.1 and glutamine synthase (GS)

Having found that pilocarpine-induced status epilepticus increases DNA methylation in astrocytes with overexpression of proinflammatory genes and downregulation of homeostatic astroglial genes; we analyzed if these alterations were sustained in the long term and thus also present in astrocytes from the epileptic brain tissue resected from drug-resistant TLE patients. For that purpose, brain sections from TLE patients and controls were processed by immunohistochemistry. As shown in Figure 5A, reactive astrocytes with increased GFAP expression, enlarged projections, and hypertrophied soma were observed in the hippocampal resected tissue from TLE patients while control brains showed typical features of human GFAP+ astrocytes (Figure 5B). Increased DNA methylation as assessed by the 5mC content in astrocytes was also observed and quantified in these brain tissue from TLE patients (Figure 5B). These samples were also analyzed for the homeostatic gene expression focusing on the Kir4.1 K^+^ rectifying channel and glutamine synthase enzyme required for the glutamate processing following astrocytic glutamate uptake. Expression of both proteins was reduced in the reactive astrocytes observed in the epileptic brain tissue resected from TLE patients (Figure 6A-B). Quantitative analysis also showed a statistically significant reduction compared to control brains (Figures 6A-B).

**Figure 5:**
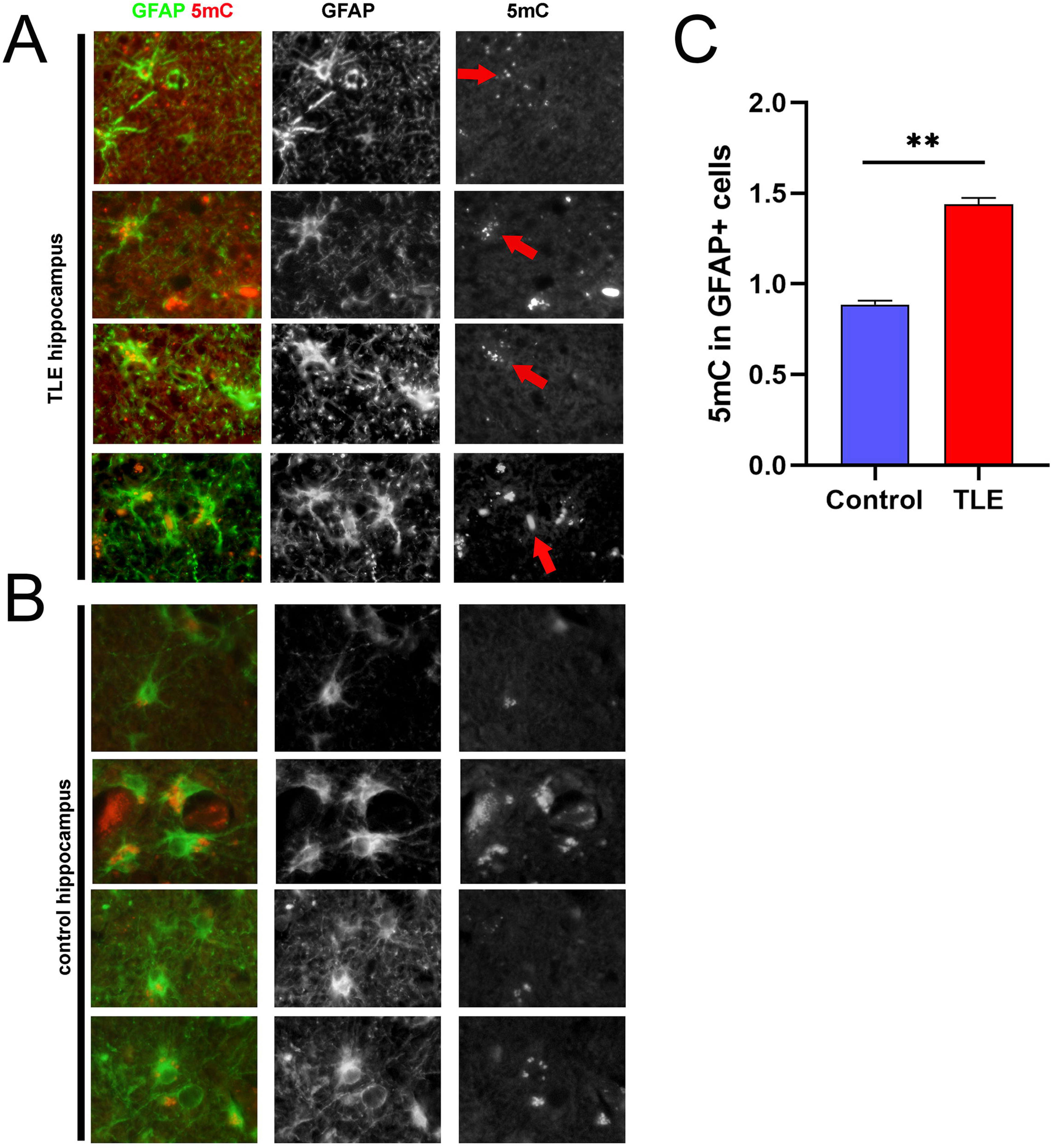
Increased DNA methylation in human brain samples from drug-resistant TLE patients. **A**: Double immunostained hippocampal brain sections form resected foci in TLE patients studied with anti-GFAP (green) and anti-5mC (red). **B**: Sections from post-mortem control material showed that the same hippocampal regions for GFAP/5mC showing a reduced degree of methylation. **C**: Quantitative analysis of the assay was presented as mean ± SEM and statistical analysis was performed with an unpaired t-test. N = 4 (TLE); N= 3 (controls).

**Figure 6:**
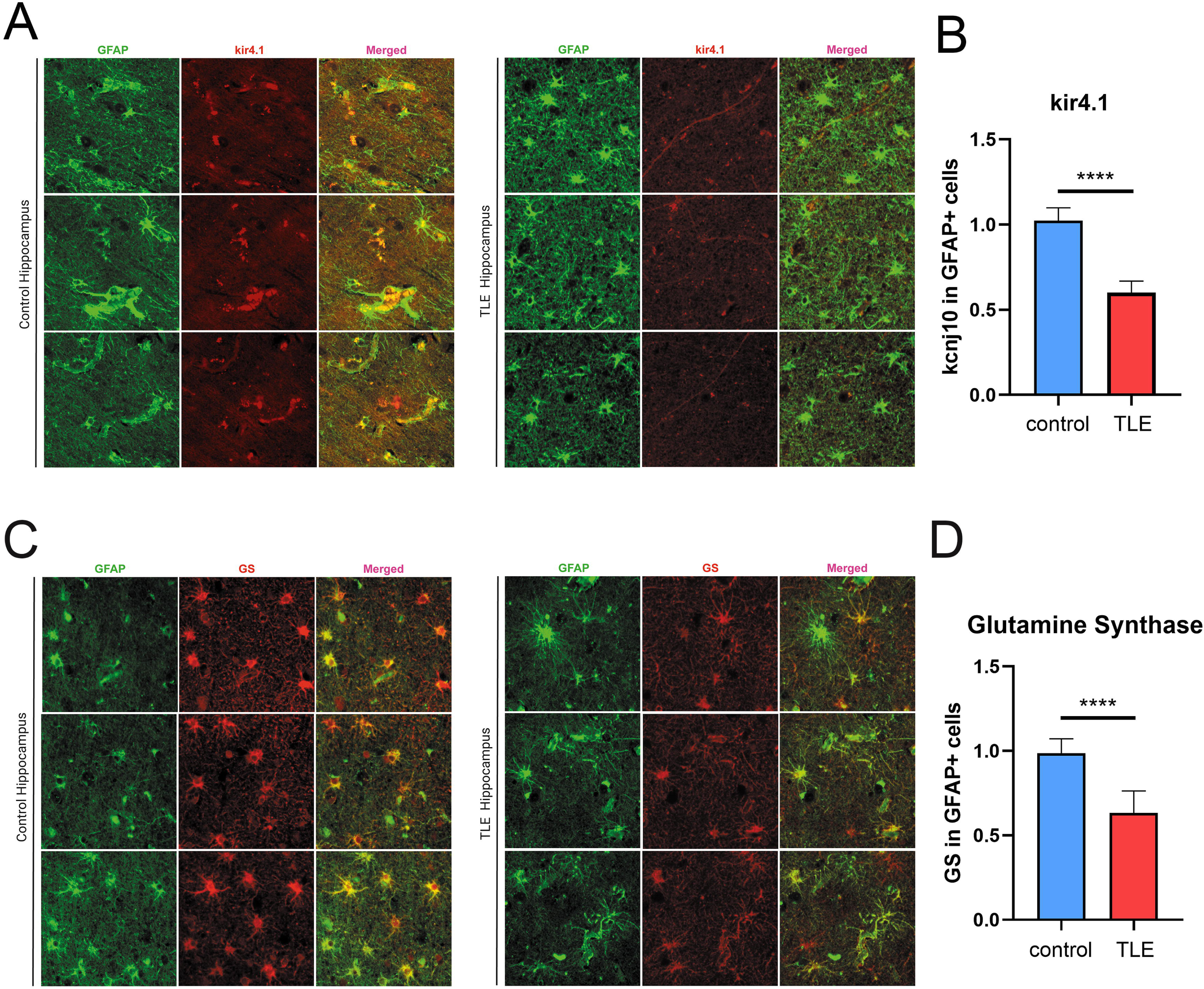
Reduced expression of homeostatic genes Kir4.1 and glutamine synthase (GS). **A**: Double immunostained hippocampal brain sections form resected foci in TLE patients or control hippocampal brain sections studied with anti-GFAP (green) and anti-Kir4.1 (red). **B**: Quantitative analysis of the assay was presented as mean ± SEM and statistical analysis was performed with an unpaired t-test. N = 4 (TLE); N= 3 (controls). **C**: Double immunostained hippocampal brain sections form resected foci in TLE patients or control hippocampal brain sections studied with anti-GFAP (green) and anti-GS (red). **D**: Quantitative analysis of the assay was presented as mean ± SEM and statistical analysis was performed with an unpaired t-test. N = 4 (TLE); N= 3 (controls).

## Discussion

The role of epigenetics in epilepsy, particularly in epileptogenesis, has been proposed, and evidence from animal models and patients has shown that epigenetic marks are altered in this pathology. Specifically, profound alterations in the level of DNA methylation in animal models and patient samples were observed in several studies (Sen et al., 2019; Tao et al., 2021; Zhang et al., 2021; Ryley Parrish et al., 2013; Bahabry et al., 2024). However, most of these studies did not account for the specific cell types in the CNS, instead analyzing global brain methylation or hydroxymethylation levels (Bahabry et al., 2024). This limitation underscores the importance of examining cell-type-specific epigenetic alterations, such as those in astrocytes.

Astrocytes are essential cells for neuronal survival in the CNS due to their metabolic coupling with neurons (Pellerin et al., 1998; Pellerin and Magistretti, 2012). In addition, astrocytes are important to modulate excitatory glutamate levels through the uptake and subsequent glutamine synthesis (Sandhu et al., 2021; Andersen and Schousboe, 2023) and modulate potassium levels via their Kir4.1 channels (Wilson, 1997; Verkhratsky and Butt, 2023). Additionally, astrocytes are pivotal for maintaining blood-brain barrier (BBB) integrity and they facilitate the extrusion of molecules (Lazarowski et al., 2004). Astrocytes are profoundly altered in epilepsy, and they are proposed as essential participants in epileptogenesis (Rossi et al., 2012, 2017). Alterations in the epigenetic pattern of DNA methylation in astrocytes were specifically observed in the Kir4.1 gene and it is proposed that this epigenetic modification facilitates neuronal excitability and epileptogenesis by impairing the K+ buffering (Boni et al., 2020; Carcak et al., 2023). Increased glutamate levels have been observed in TLE patients and experimental models and the astroglial impairment in the control of glutamate balance also seems to be dependent on epigenetic mechanisms, in this case mediated by the m6A-YTHDC2-xCT pathway (Zhang et al., 2024).

We here found that astroglial DNA becomes rapidly hypermethylated by the SE increasing their content in 5-mC, a phenomenon that occurs simultaneously with reactive astrogliosis. Interestingly, the astroglial hypermethylation persists along the analyzed time course following the pilocarpine-induced SE. In agreement with the long-term persistence of the hypermethylation, samples resected from drug-resistant TLE patients also showed an increased abundance of 5mC in reactive astrocytes, while similar age- and sex- non-TLE control patients did not show increased astroglial 5mC levels in astrocytes. Our exploration of some classical markers of proinflammatory astrocytes, specifically C3 and MAFG showed that pilocarpine-induced SE in rats facilitates the initial astroglial conversion to the proinflammatory phenotype, although the level of these proinflammatory markers tends to be normalized by 35DPSE. These results are compatible with a proinflammatory burst due to the status epilepticus that, however, is limited after several weeks. Interestingly, we observed that this proinflammatory burst is accompanied by a long-lasting repression of essential homeostatic astroglial genes Kir4.1, AQP4 and glutamine synthase in the weeks that follow pilocarpine-induced SE in experimental rats. AQP4 is the main responsible of water management in the CNS and its alterations have serious consequences for water homeostasis and edema formation in the CNS. As stated above, glutamine synthetase (GS) plays a critical role in astrocytes, acting as a key regulator of ammonia homeostasis and glutamate recycling in the brain. The long-lasting GS downregulation observed after pilocarpine-induced SE is probably contributing to the observed pro-epileptogenic glutamate disbalance in experimental models and human TLE patients (Zhang et al., 2024). In agreement with this scenario, the brain samples resected from human TLE patients also showed decreased GS expression when compared with non-disease samples. Kir4.1 expression also showed paralleled down-regulation in SE-exposed rats and brain samples resected from drug-resistant TLE patients. The consequences of impaired Kir4.1 expression are also known to be pro-epileptogenic (Boni et al., 2020; Nwaobi et al, 2014).

DNA methylation is controlled by the balance between methylation and demethylation by TET enzymes. Also, adaptor proteins belonging to the methyl-CpG binding domain family (MBD) such as MeCP2 are known to produce epileptogenic consequences when they are altered (reviewed in Gold et al., 2024). In addition, the role of de-methylation and TET enzymes in epilepsy has been recently proposed by Bahabry and colleagues (2024). DNMT1 is primarily responsible for maintaining DNA methylation during cell duplication. In contrast, DNMT3A and DNMT3B are de novo methyltransferases, which establish new methylation marks. In agreement with the hypermethylation pattern observed in astroglial DNA, we detected an early increase in the expression of DNMT3A by 7DPSE in the pilocarpine-induced SE in rats that persisted elevated until 21DPSE. DNMT1, on the other hand, showed an increase of 21DPSE and 35DPSE.

In vitro experiments provided some insights into the mechanisms of astroglial hypermethylation that seems to correlate with repression of homeostatic genes and proinflammatory burst. In fact, primary astrocytes exposed to the well-described pro-epileptogenic DAMP named HMGB1 showed long-term DNA hypermethylation as well as increased expression of proinflammatory cytokines IL-1B and IL-6, concomitantly with a 72-h peak of DNMT3A and long-lasting DNMT1 overexpression. Homeostatic genes involved in CNS homeostasis including Kir4.1 (Kcnj10), LDHA, MCT1 (Scl16a1), and GS were all downregulated in HMGB1-exposed astrocytes. Particularly lactate dehydrogenase A (LDHA) and monocarboxylate transporter 1 (MCT1) play key roles in the astroglial lactate shuttle, a mechanism essential for coupling energy metabolism between astrocytes and neurons. LDHA, predominantly expressed in astrocytes, catalyzes the conversion of pyruvate to lactate, facilitating glycolytic flux and lactate production under aerobic conditions. This lactate is then exported by MCT1, a proton-coupled monocarboxylate transporter expressed on the astrocytic membrane, to the extracellular space where it can be taken up by neurons via their MCT2 (Pellerin et al., 1998; Bergersen & Gjedde, 2012). Within neurons, lactate serves as a critical substrate for oxidative phosphorylation, supporting energy demands during synaptic activity. Disruption of LDHA or MCT1 expression is known to compromise neuronal function and has been linked to neurodegenerative conditions and cognitive deficits (Jha et al., 2015; Barros & Deitmer, 2010; Bolaños, 2016). The observed downregulation of LDHA and MCT1 expression induced by HMGB1 exposure would have neurodegenerative consequences as previously shown (Gomez Cuautle et al., 2024). Mechanistically, our methylation-sensitive studies on the GS, LDHA, and MCT1 gene promoters have shown that these promoters are hypermethylated following HMGB1 exposure.

In conclusion, our work has shown that pilocarpine-induced SE produces profuse reactive astrogliosis accompanied by long-term astroglial DNA hypermethylation, features also found in brain tissue resected from drug-resistant TLE patients. The astroglial hypermethylation correlates with the long-term downregulation of essential CNS homeostatic genes such as GS, Kir4.1 and AQP4 and a proinflammatory burst in the initial time points following SE. Mechanistically, in vitro studies showed that astroglial hypermethylation seems to be induced by the epileptogenic DAMP HMGB1 that induces the DNMT1 and DNMT3A expression and subsequent methylation of essential gene promoters in astrocytes. Taken together our findings show that astroglial DNA methylation seems to be a key step in the progression and stabilization of the epileptogenic phenotype in the brain. Strategies aimed to prevent the astroglial hypermethylation during the epileptogenic period that follows an IPE may have a profound impact in preventing the development of epileptogenesis and thus be able to modify the natural story of the disease.

## Acknowledgments

This work has been supported by grants from CONICET, UBACYT, and MinCyt (PICT 2019-0851, PICT 2017-2203 and PICT 2021-0760). The authors thank Marianella Ceol and Andrea Pecile for the animal care, Nerina Villalba for their technical assistance with microscopy facilities, and Franco Dolcetti for the support of the tissue culture room.

## References

1. Andersen JV, Schousboe A. Milestone Review: Metabolic dynamics of glutamate and GABA mediated neurotransmission - The essential roles of astrocytes. J Neurochem. 2023 Jul;166(2):109–137. doi: 10.1111/jnc.15811. Epub 2023 Mar 29.

2. Aviles-Reyes RX, Angelo MF, Villarreal A, Rios H, Lazarowski A, Ramos AJ. Intermittent hypoxia during sleep induces reactive gliosis and limited neuronal death in rats: implications for sleep apnea. J Neurochem. 2010 Feb;112(4):854–69. doi: 10.1111/j.1471-4159.2009.06535.x. Epub 2009 Dec 10.

3. Bahabry R, Hauser RM, Sánchez RG, Jago SS, Ianov L, Stuckey RJ, Parrish RR, Ver Hoef L, Lubin FD. Alterations in DNA 5-hydroxymethylation patterns in the hippocampus of an experimental model of chronic epilepsy. Neurobiol Dis. 2024 Oct 1;200:106638. doi: 10.1016/j.nbd.2024.106638. Epub 2024 Aug 13.

4. Barros, L. F., & Deitmer, J. W. (2010). Glucose and lactate supply to the synapse. Brain Research Reviews, 63(1–2), 149–159. doi10.1016/J.BRAINRESREV.2009.10.002

5. Baylin, S. B., & Jones, P. A. (2011). A decade of exploring the cancer epigenome biological and translational implications. Nature Reviews Cancer, 11(10), 726–734.

6. Berger, T. C., Vigeland, M. D., Hjorthaug, H. S., Etholm, L., Nome, C. G., Taubøll, E., Heuser, K., & Selmer, K. K. (2019). Neuronal and glial DNA methylation and gene expression changes in early epileptogenesis. PLoS ONE, 14(12). 10.1371/journal.pone.0226575

7. Bergersen LH, Gjedde A. Is lactate a volume transmitter of metabolic states of the brain? Front Neuroenergetics. 2012 Mar 19;4:5. doi: 10.3389/fnene.2012.00005.

8. Bergersen, L., & Gjedde, A. (2012). Is lactate a volume transmitter of metabolic states of the brain? Frontiers in Neuroenergetics, 4, 5.

9. Bertogliat, M. J., Morris-Blanco, K. C., & Vemuganti, R. (2020). Epigenetic mechanisms of neurodegenerative diseases and acute brain injury. Neurochemistry International, 133. 10.1016/j.neuint.2019.104642

10. Binder, D. K., et al. (2004). Astrocytes and epilepsy. Neurotherapeutics, 1(3), 406–414.

11. Bolaños, J. P. (2016). Bioenergetics and redox adaptations of astrocytes to neuronal activity. Journal of Neurochemistry, 139 Suppl 2(Suppl Suppl 2), 115–125. 10.1111/JNC.13486

12. Boni JL, Kahanovitch U, Nwaobi SE, Floyd CL, Olsen ML. DNA methylation: A mechanism for sustained alteration of KIR4.1 expression following central nervous system insult. Glia. 2020 Jul;68(7):1495–1512. doi: 10.1002/glia.23797. Epub 2020 Feb 18.

13. Çarçak N, Onat F, Sitnikova E. Astrocytes as a target for therapeutic strategies in epilepsy: current insights. Front Mol Neurosci. 2023 Jul 31;16:1183775. doi: 10.3389/fnmol.2023.1183775.

14. Cieri MB, Villarreal A, Gomez-Cuautle DD, Mailing I, Ramos AJ. Progression of reactive gliosis and astroglial phenotypic changes following stab wound-induced traumatic brain injury in mice. J Neurochem. 2023 Oct;167(2):183–203. doi: 10.1111/jnc.15941. Epub 2023 Aug 17.

15. Clossen, B. L., & Reddy, D. S. (2017). Novel therapeutic approaches for disease-modification of epileptogenesis for curing epilepsy. Biochimica et Biophysica Acta. Molecular Basis of Disease, 1863(6), 1519–1538. doi 10.1016/J.BBADIS.2017.02.003

16. D’Alessio L, Konopka H, Solís P, Scévola L, Lima MF, Nuñez C, Seoane E, Oddo S, Kochen S. Depression and Temporal Lobe Epilepsy: Expression Pattern of Calbindin Immunoreactivity in Hippocampal Dentate Gyrus of Patients Who Underwent Epilepsy Surgery with and without Comorbid Depression. Behav Neurol. 2019 May 2;2019:7396793. doi: 10.1155/2019/7396793.

17. D’Alessio L, Mesarosova L, Anink JJ, Kochen S, Solís P, Oddo S, Konopka H, Iyer AM, Mühlebner A, Lucassen PJ, Aronica E, van Vliet EA. Reduced expression of the glucocorticoid receptor in the hippocampus of patients with drug-resistant temporal lobe epilepsy and comorbid depression. Epilepsia. 2020 Aug;61(8):1595–1605. doi: 10.1111/epi.16598. Epub 2020 Jul 11.

18. Escartin C, Galea E, Lakatos A, O’Callaghan JP, et al. Reactive astrocyte nomenclature, definitions, and future directions. Nat Neurosci. 2021 Mar;24(3):312–325. doi: 10.1038/s41593-020-00783-4. Epub 2021 Feb 15.

19. Fisher, R. S., et al. (2014). ILAE Official Report: A practical clinical definition of epilepsy. Epilepsia, 55(4), 475–482.

20. Gold WA, Percy AK, Neul JL, Cobb SR, Pozzo-Miller L, Issar JK, Ben-Zeev B, Vignoli A, Kaufmann WE. Rett syndrome. Nat Rev Dis Primers. 2024 Nov 7;10(1):84. doi: 10.1038/s41572-024-00568-0.

21. Gomez-Cuautle DG, Donna S, Cieri MB, Villarreal A, Ramos AJ. Pathological remodeling of reactive astrocytes: Involvement of DNA methylation and downregulation of homeostatic genes. J Neurochem. 2024 Sep;168(9):2935–2955. doi: 10.1111/jnc.16164. Epub 2024 Jun 29.

22. Hsieh, J., & Eisch, A. J. (2010). Epigenetics, hippocampal neurogenesis, and neuropsychiatric disorders: Unraveling the genome to understand the mind. Neurobiology of Disease, 39(1), 73–84.

23. Jha, M. K., Lee, I. K., Suk, K., & Lee, W. H. (2015). Metabolic connection of inflammatory diseases: Lactate as a signaling molecule in inflammation. Trends in Neurosciences, 38(10), 634–646.

24. Kobow, K., & Blümcke, I. (2018). Epigenetics in epilepsy. Neuroscience Letters, 667, 40–46. doi 10.1016/j.neulet.2017.01.012

25. Kobow, K., et al. (2012). Epigenetic characterization of human epileptic brain tissue. Annals of Neurology, 71(4), 604–614.

26. Kwan P, Arzimanoglou A, Berg AT, Brodie MJ, Allen Hauser W, Mathern G, Moshé SL, Perucca E, Wiebe S, French J. Definition of drug resistant epilepsy: consensus proposal by the ad hoc Task Force of the ILAE Commission on Therapeutic Strategies. Epilepsia. 2010 Jun;51(6):1069–77. doi: 10.1111/j.1528-1167.2009.02397.x.

27. Kwan, P., & Brodie, M. J. (2000). Early identification of refractory epilepsy. New England Journal of Medicine, 342(5), 314–319.

28. Lazarowski A, Czornyj L. Potential role of multidrug resistant proteins in refractory epilepsy and antiepileptic drugs interactions. Drug Metabol Drug Interact. 2011;26(1):21–6. doi: 10.1515/DMDI.

29. Lazarowski A, Ramos AJ, García-Rivello H, Brusco A, Girardi E. Neuronal and glial expression of the multidrug resistance gene product in an experimental epilepsy model. Cell Mol Neurobiol. 2004 Feb;24(1):77–85. doi: 10.1023/b:cemn.0000012726.43842.d2.

30. Li, E., Bestor, T. H., & Jaenisch, R. (1992). Targeted mutation of the DNA methyltransferase gene results in embryonic lethality. Cell, 69(6), 915–926.

31. Łukasiuk K, Lasoń W. Emerging Molecular Targets for Anti-Epileptogenic and Epilepsy Modifying Drugs. Int J Mol Sci. 2023 Feb 2;24(3):2928. doi: 10.3390/ijms24032928.

32. Maroso M, Balosso S, Ravizza T, Liu J, Aronica E, Iyer AM, Rossetti C, Molteni M, Casalgrandi M, Manfredi AA, Bianchi ME, Vezzani A. Toll-like receptor 4 and high-mobility group box-1 are involved in ictogenesis and can be targeted to reduce seizures. Nat Med. 2010 Apr;16(4):413–9. doi: 10.1038/nm.2127.

33. Martins-Ferreira R, Leal B, Chaves J, Ciudad L, Samões R, Martins da Silva A, Pinho Costa P, Ballestar E. Circulating cell-free DNA methylation mirrors alterations in cerebral patterns in epilepsy. Clin Epigenetics. 2022 Dec 28;14(1):188. doi: 10.1186/s13148-022-01416-2.

34. McNamara JO, Huang YZ, Leonard AS. Molecular signaling mechanisms underlying epileptogenesis. Sci STKE. 2006 Oct 10;2006(356):re12. doi: 10.1126/stke.3562006re12.

35. Miller-Delaney, S. F., et al. (2015). Differential DNA methylation profiles of coding and non-coding genes define hippocampal sclerosis in human temporal lobe epilepsy. Brain, 138(3), 616–631.

36. Nwaobi SE, Lin E, Peramsetty SR, Olsen ML. DNA methylation functions as a critical regulator of Kir4.1 expression during CNS development. Glia. 2014 Mar;62(3):411–27. doi: 10.1002/glia.22613. Epub 2014 Jan 10.

37. Okano, M., Bell, D. W., Haber, D. A., & Li, E. (1999). DNA methyltransferases Dnmt3a and Dnmt3b are essential for de novo methylation and mammalian development. Cell, 99(3), 247–257.

38. Pellerin L, Magistretti PJ. Sweet sixteen for ANLS. J Cereb Blood Flow Metab. 2012 Jul;32(7):1152–66. doi: 10.1038/jcbfm.2011.149. Epub 2011 Oct 26.

39. Pellerin, L., Pellegri, G., Martin, J. L., & Magistretti, P. J. (1998). Expression of monocarboxylate transporter MCT1 in the central nervous system: a marker of the astrocyte–neuron lactate shuttle. Proceedings of the National Academy of Sciences, 95(11), 6065–6070.

40. Purnell, B. S., Alves, M., & Boison, D. (2023). Astrocyte-neuron circuits in epilepsy. Neurobiology of Disease, 179, 106058. 10.1016/j.nbd.2023.106058

41. Ravizza T, Terrone G, Salamone A, Frigerio F, Balosso S, Antoine DJ, Vezzani A. High Mobility Group Box 1 is a novel pathogenic factor and a mechanistic biomarker for epilepsy. Brain Behav Immun. 2018 Aug;72:14–21. doi: 10.1016/j.bbi.2017.10.008.

42. Reddy, S. D., Clossen, B. L., & Reddy, D. S. (2018). Epigenetic Histone Deacetylation Inhibition Prevents the Development and Persistence of Temporal Lobe Epilepsy. The Journal of Pharmacology and Experimental Therapeutics, 364(1), 97–109. 10.1124/JPET.117.244939

43. Robel, S., et al. (2015). Epigenomic and transcriptomic changes during astrocyte activation by elevated extra cellular K+. Journal of Neuroscience, 35(2), 634–650.

44. Rosciszewski G, Cadena V, Auzmendi J, Cieri MB, Lukin J, Rossi AR, Murta V, Villarreal A, Reinés A, Gomes FCA, Ramos AJ. Detrimental Effects of HMGB-1 Require Microglial-Astroglial Interaction: Implications for the Status Epilepticus -Induced Neuroinflammation. Front Cell Neurosci. 2019 Aug 27;13:380. doi: 10.3389/fncel.2019.00380.

45. Rosciszewski G, Cadena V, Murta V, Lukin J, Villarreal A, Roger T, Ramos AJ. Toll-Like Receptor 4 (TLR4) and Triggering Receptor Expressed on Myeloid Cells-2 (TREM-2) Activation Balance Astrocyte Polarization into a Proinflammatory Phenotype. Mol Neurobiol. 2018 May;55(5):3875–3888. doi: 10.1007/s12035-017-0618-z. Epub 2017 May 25.

46. Rossi A, Murta V, Auzmendi J, Ramos AJ. Early Gabapentin Treatment during the Latency Period Increases Convulsive Threshold, Reduces Microglial Activation and Macrophage Infiltration in the Lithium-Pilocarpine Model of Epilepsy. Pharmaceuticals (Basel). 2017 Nov 28;10(4):93. doi: 10.3390/ph10040093.

47. Rossi AR, Angelo MF, Villarreal A, Lukin J, Ramos AJ. Gabapentin administration reduces reactive gliosis and neurodegeneration after pilocarpine-induced status epilepticus. PLoS One. 2013 Nov 8;8(11):e78516. doi: 10.1371/journal.pone.0078516.

48. Ryley Parrish R, Albertson AJ, Buckingham SC, Hablitz JJ, Mascia KL, Davis Haselden W, Lubin FD. Status epilepticus triggers early and late alterations in brain-derived neurotrophic factor and NMDA glutamate receptor Grin2b DNA methylation levels in the hippocampus. Neuroscience. 2013 Sep 17;248:602–19. doi: 10.1016/j.neuroscience.2013.06.029. Epub 2013 Jun 27.

49. Sandhu MRS, Gruenbaum BF, Gruenbaum SE, Dhaher R, Deshpande K, Funaro MC, Lee TW, Zaveri HP, Eid T. Astroglial Glutamine Synthetase and the Pathogenesis of Mesial Temporal Lobe Epilepsy. Front Neurol. 2021 Apr 13;12:665334. doi: 10.3389/fneur.2021.665334.

50. Sarchi PV, Gomez Cuautle D, Rossi A, Ramos AJ. Participation of the spleen in the neuroinflammation after pilocarpine-induced status epilepticus: implications for epileptogenesis and epilepsy. Clin Sci (Lond). 2024 May 8;138(9):555–572. doi: 10.1042/CS20231621.

51. Sen K, Gadkari R, Agarwal R, Sundaram S. Differential DNA Methylation Patterns in Patients with Epilepsy due to Malformations of Cortical Development: A Pilot Study. Neurol India. 2019 Nov-Dec;67(6):1469–1471. doi: 10.4103/0028-3886.273638.

52. Tao H, Chen Z, Wu J, Chen J, Chen Y, Fu J, Sun C, Zhou H, Zhong W, Zhou X, Li K. DNA Methylation Signature of Epileptic Encephalopathy-Related Pathogenic Genes Encoding Ion Channels in Temporal Lobe Epilepsy. Front Neurol. 2021 Jul 29;12:692412. doi: 10.3389/fneur.2021.692412.

53. Vargas-Sánchez, K., et al. (2019). Reactive astrocytes in epilepsy: Friends or foes? Current Neuropharmacology, 17(10), 922–942.

54. Verkhratsky A, Butt AM. Neuroglia: Function and Pathology. Elsevier; 2023:713.

55. Verkhratsky A, Parpura V, Li B, Scuderi C. Astrocytes: The Housekeepers and Guardians of the CNS. Adv Neurobiol. 2021;26:21–53. doi: 10.1007/978-3-030-77375-5_2.

56. Verkhratsky A, Zorec R, Rodriguez JJ, Parpura V. Neuroglia: Functional Paralysis and Reactivity in Alzheimer’s Disease and Other Neurodegenerative Pathologies. Adv Neurobiol. 2017;15:427–449. doi: 10.1007/978-3-319-57193-5_17.

57. Vezzani A, Ravizza T, Bedner P, Aronica E, Steinhäuser C, Boison D. Astrocytes in the initiation and progression of epilepsy. Nat Rev Neurol. 2022 Dec;18(12):707–722. doi: 10.1038/s41582-022-00727-5. Epub 2022 Oct 24.

58. Villarreal A, Vidos C, Monteverde Busso M, Cieri MB, Ramos AJ. Pathological Neuroinflammatory Conversion of Reactive Astrocytes Is Induced by Microglia and Involves Chromatin Remodeling. Front Pharmacol. 2021 Jun 21;12:689346. doi: 10.3389/fphar.2021.689346.

59. Wheeler MA, Clark IC, Tjon EC, Li Z, Zandee SEJ, Couturier CP, Watson BR, Scalisi G, Alkwai S, Rothhammer V, Rotem A, Heyman JA, Thaploo S, Sanmarco LM, Ragoussis J, Weitz DA, Petrecca K, Moffitt JR, Becher B, Antel JP, Prat A, Quintana FJ. MAFG-driven astrocytes promote CNS inflammation. Nature. 2020 Feb;578(7796):593–599. doi: 10.1038/s41586-020-1999-0. Epub 2020 Feb 12.

60. Williams-Karnesky RL, Sandau US, Lusardi TA, Lytle NK, Farrell JM, Pritchard EM, Kaplan DL, Boison D. Epigenetic changes induced by adenosine augmentation therapy prevent epileptogenesis. J Clin Invest. 2013 Aug;123(8):3552–63. doi: 10.1172/JCI65636. Epub 2013 Jul 25.

61. Wilson JX. Antioxidant defense of the brain: a role for astrocytes. Can J Physiol Pharmacol. 1997 Oct-Nov;75(10-11):1149–63.

62. Zhang, K., Yang, Z., Yang, Z., Du, L., Zhou, Y., Fu, S., Wang, X., Li, X., Liu, D., & He, X. (2024). The m6A reader YTHDC2 promotes the pathophysiology of temporal lobe epilepsy by modulating SLC7A11-dependent glutamate dysregulation in astrocytes. Theranostics, 14(14), 5551. 10.7150/thno.100703

